# Analysis and Prediction of Unplanned Intensive Care Unit Readmission using Recurrent Neural Networks with Long Short-Term Memory

**DOI:** 10.1101/385518

**Authors:** Yu-Wei Lin, Yuqian Zhou, Faraz Faghri, Michael J. Shaw, Roy H. Campbell

**Author notes:** These authors contributed equally to this work (Yu-Wei Lin, Yuqian Zhou).

## Abstract

**Background:** Unplanned readmission of a hospitalized patient is an extremely undesirable outcome as the patient may have been exposed to additional risks. The rates of unplanned readmission are, therefore, regarded as an important performance indicator for the medical quality of a hospital and healthcare system. Identifying high-risk patients likely to suffer from readmission before release benefits both the patients and the medical providers. The emergence of machine learning to detect hidden patterns in complex, multi-dimensional datasets provides unparalleled opportunities to develop efficient discharge decision-making support system for physicians.

**Methods and Findings:** We used supervised machine learning approaches for ICU readmission prediction. We used machine learning methods on comprehensive, longitudinal clinical data from the MIMIC-III to predict the ICU readmission of patients within 30 days of their discharge. We have utilized recent machine learning techniques such as Recurrent Neural Networks (RNN) with Long Short-Term Memory (LSTM), by this we have been able incorporate the multivariate features of EHRs and capture sudden fluctuations in chart event features (e.g. glucose and heart rate) that are significant in time series with temporal dependencies, which cannot be properly captured by traditional static models, but can be captured by our proposed deep neural network based model. We incorporate multiple types of features including chart events, demographic, and ICD9 embeddings. Our machine learning models identifies ICU readmissions at a higher sensitivity rate (0.742) and an improved Area Under the Curve (0.791) compared with traditional methods. We also illustrate the importance of each portion of the features and different combinations of the models to verify the effectiveness of the proposed model.

**Conclusion:** Our manuscript highlights the ability of machine learning models to improve our ICU decision making accuracy, and is a real-world example of precision medicine in hospitals. These data-driven results enable clinicians to make assisted decisions within their patient cohorts. This knowledge could have immediate implications for hospitals by improving the detection of possible readmission. We anticipate that machine learning models will improve patient counseling, hospital administration, allocation of healthcare resources and ultimately individualized clinical care.

## Introduction

Unplanned hospital readmission is an avoidable waste of medical resources and indicates that patients may have been placed at risk. Therefore, to solve this problem, in 2010, the Affordable Care Act (ACA) created the Hospital Readmissions Reduction Program to penalize the hospitals whose 30-day readmission rates are higher than expected [1]. The financial penalties are given by the Centers for Medicare & Medicaid Services by reducing payments to hospitals [2].

In addition to hospital readmission, intensive care unit (ICU) readmission brings further financial risk, morbidity and mortality risks [3, 4]. Reported by Kaben et al 2008, the mortality rates of ICU readmitted patients ranges approximately from 26 % to 58 % [5]. That is, premature ward-level care transition or discharge from ICU exposes patients to the risks of unsuitable treatment, which further leads to an avoidable mortality [6]. Surprisingly, even in developed countries, hospitals suffer from high ICU readmission rates, around 10 % patients will be readmitted back to ICU within a hospital stay [3]. Moreover, as reported by Kramer et al. 2013, in the U.S. there is an escalating trend for ICU readmission rates rising from 4.6 % in 1989 to 6.4 % in 2003 [4]. ICU readmission rates, therefore, become one of the critical quality indicators in ICU performance evaluation.

According to the recent studies, 27 % to 42 % of ICU readmitted patients are discharged from ICU prematurely [3, 7]. To reduce avoidable ICU readmission, we need to identify patients with a higher risk of ICU readmission [8]. Is this way, the physicians can relocate the additional medical resources for care delivery used in unncessary readmission to put more emphasis on patients with greater needs. Avoiding unnecessary readmission is even more important in the ICU than in the general hospital because ICU resources are relatively scarce. Consequently, an efficient discharge decision-making support system that can assist ICU physicians to identify more accurately those patients with a high risk of hospital readmission would be beneficial.

Data-driven predictive models aimed at predicting ICU readmission may be built from many different data sets including administrative claims [9–11], insurance claims, and electronic health records (EHRs). Insurance claims models are not practical for real-time prediction [12]. Electronic health records (EHR) have proved to provide appropriate data for medical strategy design support. A systematic review of readmission prediction models [13], Kansagara et al. summarize 26 unique readmission prediction models of which 23 models rely on EHR. Therefore, the most recent work focuses on predicting all-cause 30-day readmission using EHR data. Jamei et al. [12] proposed an accurate and real-time prediction model based on neural networks trained from EHR data.

Even though previous predictive models have been studied to resolve the problem of identifying patients with a high-risk of readmission, these studies have drawbacks. First, the scope of the models is limited in that they are only designed for a specific disease. Most early work focuses on patients with specific diseases like heart failure [14], HIV [15], and diabetes [16], or treatments like kidney transplants [17]. Second, no model proposed in the literature has been able to predict ICU readmissions to a satisfactory degree yet [18] and the models suffer from low sensitivity of around 0.65 [6, 12, 18]. Third, the models do not exploit the time series features of EHR into consideration. Intuitively, EHR data has a sequential data structure [19]. Yang et al. embody the time series variables (e.g. average number of days between admissions and number of previous admissions) in their model, but do not consider the sequential data structures, which might lead to information loss [20]. Finally, explaining the reliability and robustness of the model is necessary for clinical applications. Few of the works attempt to understand and interpret the predictive model, especially the approaches that build “black-box” like neural networks.

In this study, we focus on the analysis and prediction of unplanned ICU readmission based on time series data. We propose a recurrent neural network (RNN) architecture with long short-term memory (LSTM) layers to learn a better predictive model that incorporates time-series. We also incorporate low-dimensional representations (embeddings) of medical concepts (e.g. diseases (ICD-9), treatment, laboratory events, etc.) as input of the model [9, 21]. Finally, we test, validate and explain the proposed methods via MIMIC-III dataset [22], containing more than 40,000 patients information and 60,000 ICU admissions records over a 10 year period [22]. We leverage the dataset to provide clinicians with data-driven decision-making support that can help prevent inappropriate discharge or transfer of patients that are high-risk for readmission so that ICU can reduce effectively the risk to the patient of readmission and reduce cost.

## Methods

To accompany this report, and to allow independent replication and extension of our work, we have made the code publicly available for use by non-profit academic researchers (https://github.com/Jeffreylin0925/MIMIC-III_ICU_Readmission_Analysis). The code is part of the supplemental information; it includes the step-by-step instructions of the statistical and machine learning analysis.

### Dataset Construction

The readmission dataset is constructed from the MIMIC-III Critical Care Database. MIMIC-III consists of the health-related EHR data of more than 40,000 patients in the Intensive Care Units (ICU) of the Beth Israel Deaconess Medical Center between 2001 and 2012. One patient may have multiple in-hospital records in the dataset. Following the data screening process stated in [19], we first screen out the patients under age 18, and remove the patients who died in the ICU. This results in totally 35,334 patients with 48,393 ICU stays. We then split the processed patients into training(80%), validation(10%)and testing(10%) partitions and conduct a five-fold cross validation.

Note that one patient may have multiple records, so the number of items may not equal in each fold.

To construct the dataset for ICU readmission, we categorize all selected patients and the corresponding ICU stay records into positive or negative cases. Specifically, the following cases are considered to be positive patient stays:

- 35,55 records: the patients were transferred to low-level wards from ICU, but returned to ICU again,
- 1,974 records: the patients were transferred to low-level wards from ICU, and died later,
- 3,205 records: the patients were discharged, but returned to the ICU within the next 30 days,
- 2,556 records: the patients were discharged, and died within the next 30 days.

Positive cases are regarded as ones in which the patients could benefit from a prediction of readmission before being transferred or discharged. Negative cases, on the contrast, are those that the patient do not need ICU readmission. Specifically, patients who were transferred or discharged from ICU and did not return and are still alive within the next 30 days are considered to be negative cases.

### Feature Extraction

In this section, we introduce the features we use in ICU Readmission prediction tasks. There are several significant groups of variables for predicting readmission. The first group of variables are chart events. Chart events are recorded from notes of health care providers (e.g., physicians and nurses) and represent the patients’ physiological conditions from experts’ observation and opinions [18]. Second, patient variables, especially chronic diseases, that are found strongly associated with ICU readmission risk [6, 23]. Thirdly, the basic demographic information, such as gender, age, race, that are again demonstrated as important factors in the state of art readmission prediction [12].

In this study, we leverage all of the above-mentioned features and further consider the time series information for the Readmission prediction tasks. Our features consist of chart events, ICD9 embeddings, and demographic information of the patients. To compare the proposed model with some traditional methods like logistic regression, we also extract statistical features from the chart events for usage.

### Chart Events

We extract 17 types of time series chart events within a 48-hour window from the MIMIC-III dataset. The raw features include numerical ones like diastolic blood pressure, and categorical items like capillary refill rate. The detailed feature items and their dimensions are shown in Table 1. We use the median from Wikipedia as the normal value for each chart event. The total dimension of the raw features from the chart events is 59. However, in the raw data for the chart events there are a few records that are missing. To identify the missing positions, we create a 17-dim binary indicator feature and append it to the chart events feature. This feature indicates whether the record of each type of chart event exists. Therefore, the total dimension of chart event feature is 48 × 76.

**Table 1.** 17 Types of Features in Chart Events

| Chart Events | Dim | Normal | Chart Events | Dim | Normal |
| --- | --- | --- | --- | --- | --- |
| 1. Glasgow coma scale eye opening | 8 | 4 Spontaneously | 11. Fraction inspired oxygen | 1 | 0.21 |
| 2. Glasgow coma scale verbal response | 12 | 5 Oriented | 12. Oxygen saturation | 1 | 97.5 |
| 3. Glasgow coma scale motor response | 12 | 6 Obeys Commands | 13. Respiratory rate | 1 | 15.0 |
| 4. Glasgow coma scale total | 13 | 15 | 14. Body Temperature | 1 | 37.0 |
| 5. Capillary refill rate | 2 | Normal <3 secs | 15. pH | 1 | 7.4 |
| 6. Diastolic blood pressure | 1 | 70.0 | 16. Weight | 1 | 80.7 |
| 7. Systolic blood pressure | 1 | 105.0 | 17. Height | 1 | 168.8 |
| 8. Mean blood pressure | 1 | 87.5 |  |  |  |
| 9. Heart Rate | 1 | 80.0 |  |  |  |
| 10. Glucose | 1 | 85.0 |  |  |  |

\*DT: Data Type, Dim: Dimension, Normal: Normal Value

### Demographic features

The demographic features we consider consist of the patients’ gender, age, race, and insurance type. The detailed categories and dimensions are summarized in table 2. The reason why we include the insurance type is in case it influences the discharge/transfer rate. For example, although unlikely, an insurance type (uninsured) could lead to insufficient payment and might result in an unexpected discharge. The whole dimension of the demographic features is 14.

**Table 2.** Demographic Features

| Chart Events | Dim | Option | Chart Events | Dim | Option |
| --- | --- | --- | --- | --- | --- |
| 1. Gender | 2 | Male/Female | 2. Age | 1 | 18-120 |
| 3. Insurance Type | 5 | Government, Self, Medicare, Private, Medicaid | 4. Race | 6 | Asian, Black, Hispanic, White, Other, No Information |

\*Dim: Dimension

### ICD9 Embeddings

In [23], Brown et al (2013) found that chronic diseases are one of the most important factors associated with later readmissions. However, the disease information in a EHR dataset are generally sparse, which makes them a poor foundation for deep learning methods.

To deal with the EHR disease data sparsity, we apply the approach presented in [9] to compute the pretrained 300-dim embedding for each ICD9 code recorded for patients. Utilizing a lower dimension embedding of ICD9 will benefit the training by avoiding a sparse representation and applying the information of the relationships among different diseases. For a patient with multiple diseases, we simply take addition of embeddings of all the diseases to form the feature.

### Times Series Window

For temporal information modeling of the time-series ICU records, we apply a 48-hour window for each ICU stay. Specifically, we claim that the data during the last 48 hours before the patient is discharged or transferred are the most informative for readmission prediction. Therefore, we use only the last 48-hour data from each ICU record. To cope with the data missing problem when the length of the record is shorter than 48 hours, we simply replicate the data of the last hour to fill the length gap.

### Statistical Features

For the implementation of the traditional methods, we also extract the statistical features within each 48-hour window. For the numerical chart events, we regress the 48 data points linearly and record the rate and the bias in the linear function. For example, after computing the linear function *y* = *ax* + *b*, we keep the 2-dim features [a,b] as the statistical information of this event. For each categorical event, such as capillary refill rate, we simply use the majority category to represent it. One example of the computation of the statistical features is illustrated in Fig 1. After computing the statistical features, each 48-hour data window will become a single data point, and the dimension of the chart events becomes 71.

**Fig 1.**
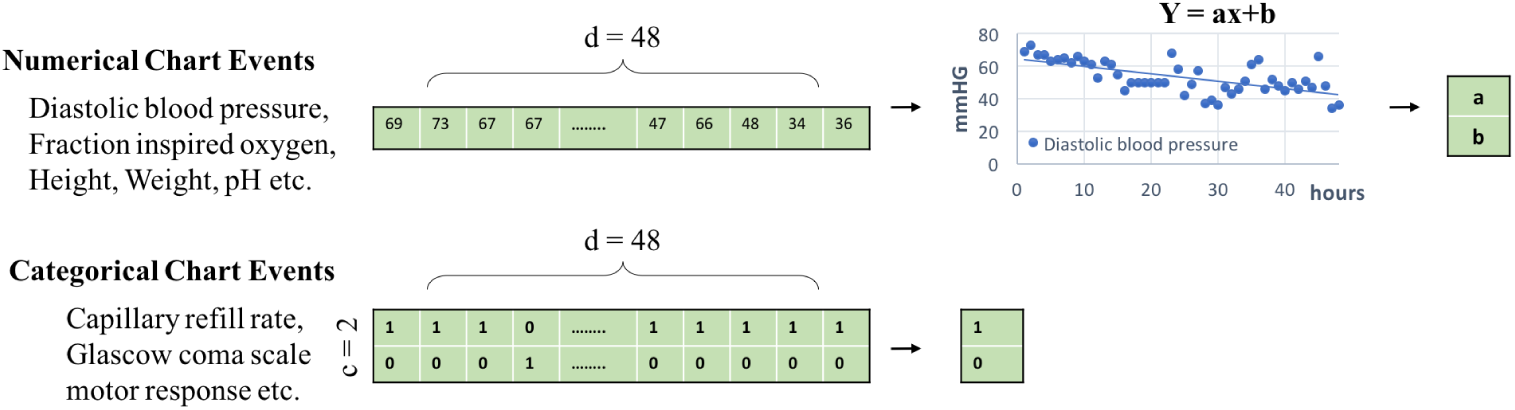
Statistical feature computation. For numerical chart events, we conduct linear regression on the 48-hour data points, and record the rate and bias value as the feature. For categorical events, we simply compute the average occurrence of the categories.

### Model Structure

Fig 2 shows the model structure of our system. As shown in Fig 2a, we utilize a bidirectional LSTM combined with an additional LSTM layer, followed by a dense decision layer with one output neuron activated by a sigmoid function. The hidden units of the LSTM layer is 16. Bidirectional LSTM learns the temporal information across the whole training window. In this case, given an ICU stay record with length *T* = 48, where the observation of each hour is denoted by *x*_*t*_ ∈ *R*^1×*D*^, and D is the feature dimension. The output of a single LSTM cell can be computed by the following equations,

**Fig 2.**
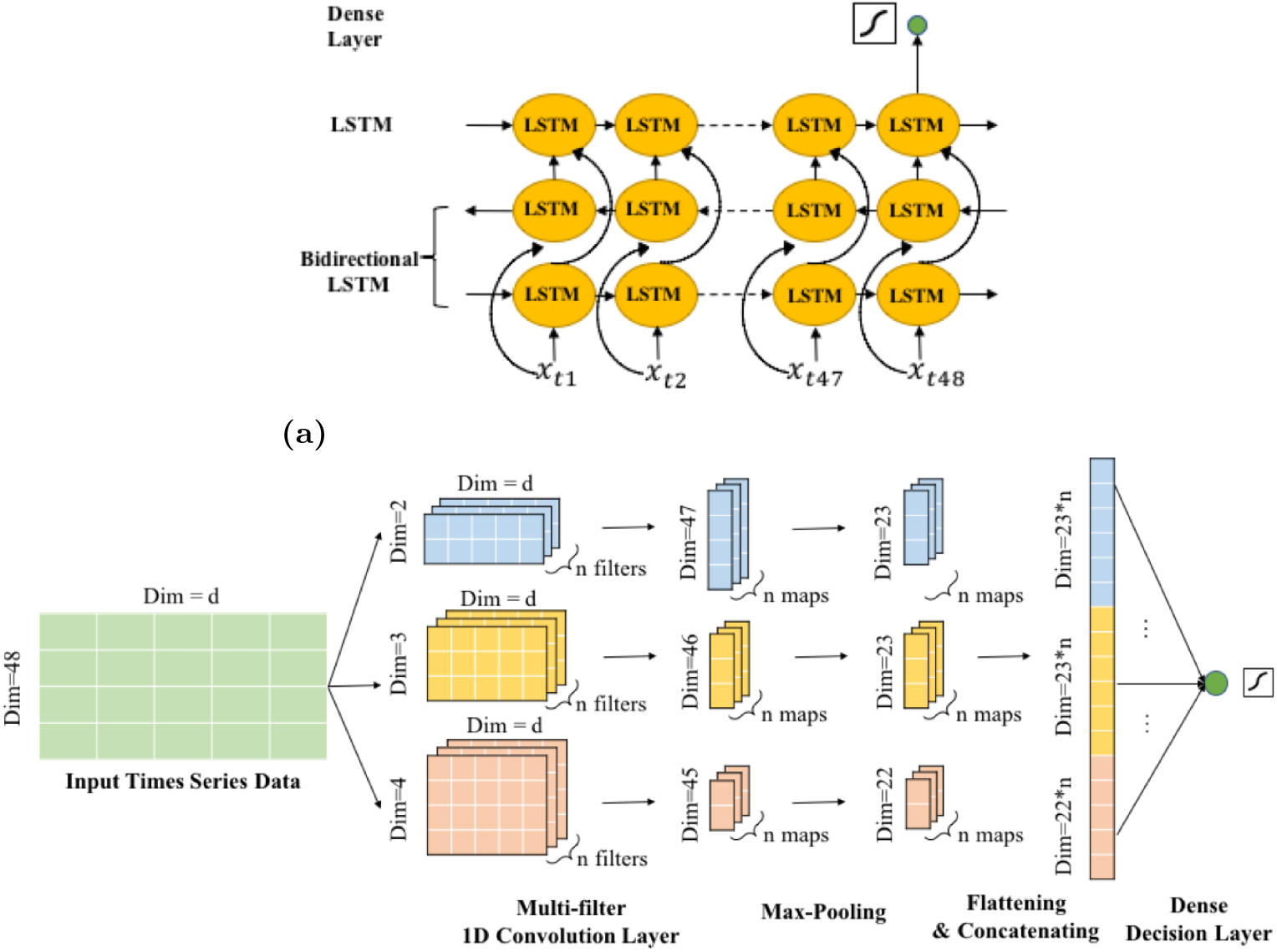
(a)A bidirectional LSTM combined with an additional LSTM layer, followed by a dense decision layer with one output neuron activated by a sigmoid function. The hidden units of the LSTM layer is 16. (b)1D Multi-filter Convolutional Neural Network. Given a 48-hour data window of dimension *D*, we conduct the convolution on the time axis with filter size 2,3 or 4. The computed feature maps are finally concatenated and fully connected to a dense decision layer with one output neuron.

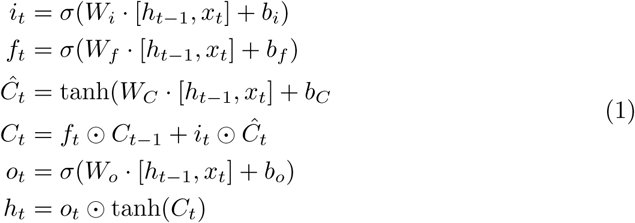

The above functions can be simply denoted by *h_t_* = *LST M* (*h_t−_*_1_*, x_t_*). We utilized the hidden value of the last time stamp to predict the readmission possibility, thus the final output after going through the dense layer would be,

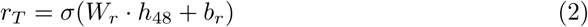

where σ is the indicator of the sigmoid activation function, and the *r*_*T*_ represents the prediction of whether this patient with the ICU stay record will have a readmission, ranging from zero to one. The dimension of *h_t_* is *R*^1×16^, so the *W*_*r*_ ∈ *R*^16×1^. We use binary cross entropy loss to update the weights.

In additional to a LSTM-based model, we also apply a CNN-based model for exploration as shown in Fig 2b. The CNN structure we build is a multi-filter CNN introduced in [24]. Given a 48-hour data window of dimension *D*, we conduct the convolution on the time axis with filter size 2,3 or 4. The computed feature maps are finally concatenated and fully connected to a dense decision layer with one output neuron. We also compare the performance of the combination of LSTM and CNN models, and the details will be introduced in the next section.

We evaluate the performance of the models by mainly using positive case recall rate (sensitivity) and Area-under-curve of ROC. The reason why this two metrics are important is that, first, recall rate of positive cases plays a more important role in screening patients. In other words, a highly sensitive test indicate that the model can correctly identifies patients with a high risk of readmission. Secondly, AUC under ROC measures the overall performance of the recall with respect to different false positive rate. Models with higher AUC under ROC will demonstrate a more powerful screening capability, benefiting the initial selection of the patient candidates for the physicians.

## Results

In this section, we illustrate the experiments we conducted to evaluate the performance of the model. It consists of traditional statistical approaches like logistic regression, random forest etc., and deep learning based temporal model like LSTM. We compared the performance obtained by different models and derived the optimal solution of the prediction system. All the models were reimplemented using keras based on the benchmark code of [19]. The learning rate of training was set to 1*e*^−3^, and we used Adam optimizer to train the model with beta 0.9. Based on the logic in [25], we trained at most 50 epochs and selected the model with the highest AUC under ROC on the validation partition. During evaluation, we set up the decision threshold as 0.5.

### Statistical Model

We trained four statistical models as our baseline, including Logistic Regression, Naive Bayes, Random Forest and SVM. The feature we utilized is illustrated in section. The results are shown in table 3. SVM outperforms other traditional methods in terms of positive recall and AUC under ROC curve, and Logistic Regression is also suitable for this readmission dataset.

**Table 3.** Performance Comparison of Different Models and Features

| Model | Feature | Acc | Pre-0 | Pre-1 | Re-0 | Re-1 | A.R | A.P |
| --- | --- | --- | --- | --- | --- | --- | --- | --- |
| **Baseline** |  |  |  |  |  |  |  |  |
| LR | L48-h STAT + ICD9 | 0.723 | 0.902 | 0.382 | 0.736 | 0.670 | 0.770 | 0.465 |
| NB | L48-h STAT + ICD9 | 0.744 | 0.859 | 0.373 | 0.814 | 0.453 | 0.709 | 0.458 |
| RF | L48-h STAT + ICD9 | 0.705 | 0.874 | 0.345 | 0.739 | 0.563 | 0.714 | 0.376 |
| SVM | L48-h STAT + ICD9 | 0.711 | 0.907 | 0.374 | 0.714 | **0.701** | **0.775** | 0.456 |
| **Feature Selection** |  |  |  |  |  |  |  |  |
| LSTM | F48-h CE + ICD9 | 0.687 | 0.912 | 0.355 | 0.676 | 0.731 | 0.777 | 0.481 |
| LSTM | L48-h CE + ICD9 | 0.707 | 0.911 | 0.372 | 0.704 | 0.717 | 0.784 | 0.504 |
| LSTM | L48-h CE | 0.676 | 0.876 | 0.325 | 0.697 | 0.593 | 0.704 | 0.406 |
| LSTM | L48-h CE + ICD9 +D | 0.695 | 0.914 | 0.363 | 0.686 | **0.733** | **0.787** | 0.509 |
| **Model Selection** |  |  |  |  |  |  |  |  |
| CNN | L48-h CE + ICD9 | 0.731 | 0.902 | 0.395 | 0.747 | 0.665 | 0.780 | 0.490 |
| CNN+LSTM | L48-h CE + ICD9 | 0.688 | 0.915 | 0.361 | 0.676 | 0.739 | 0.785 | 0.492 |
| CNN+LSTM | L48-h CE + ICD9 + D | 0.710 | 0.910 | 0.376 | 0.710 | 0.710 | 0.787 | 0.496 |
| LSTM+CNN | L48-h CE + ICD9 | 0.698 | 0.914 | 0.369 | 0.690 | 0.729 | 0.786 | 0.510 |
| LSTM+CNN | L48-h CE + ICD9 + D | 0.698 | 0.916 | 0.367 | 0.687 | **0.742** | **0.791** | 0.513 |

\*Acc:Accuracy Pre:Precision Re:Recall A.R: AUC under ROC A.P:AUC under PRC L48:Last 48 hours F48:First 48 hours CE:chart events STAT:statistical features D:Demographic features

### Feature Selection

We conduct a feature ablation test to evaluate the influence of different portions of features on the system performance. Specifically, we selected the Bidirectional LSTM as our base model, and deployed different combinations of the feature input. As shown in table 3, our results demonstrated that the last-48h features performs relatively better than the first-48h data in terms of positive case recall rate and AUC under ROC. In addition, ICD9 embedding is necessary in predicting the readmission rate. Demographic features will also greatly benefit the performance. Therefore, we claim that the full set of features involving Last-48h chart events and their identifiers, ICD9 embeddings, and demographic information perform the best among all the combinations.

### Model Selection

We attempted multiple model structures including bidirectional LSTM, CNN, and the combinations of them. The detailed combination strategies are illustrated in Fig 4.

**Fig 3.**
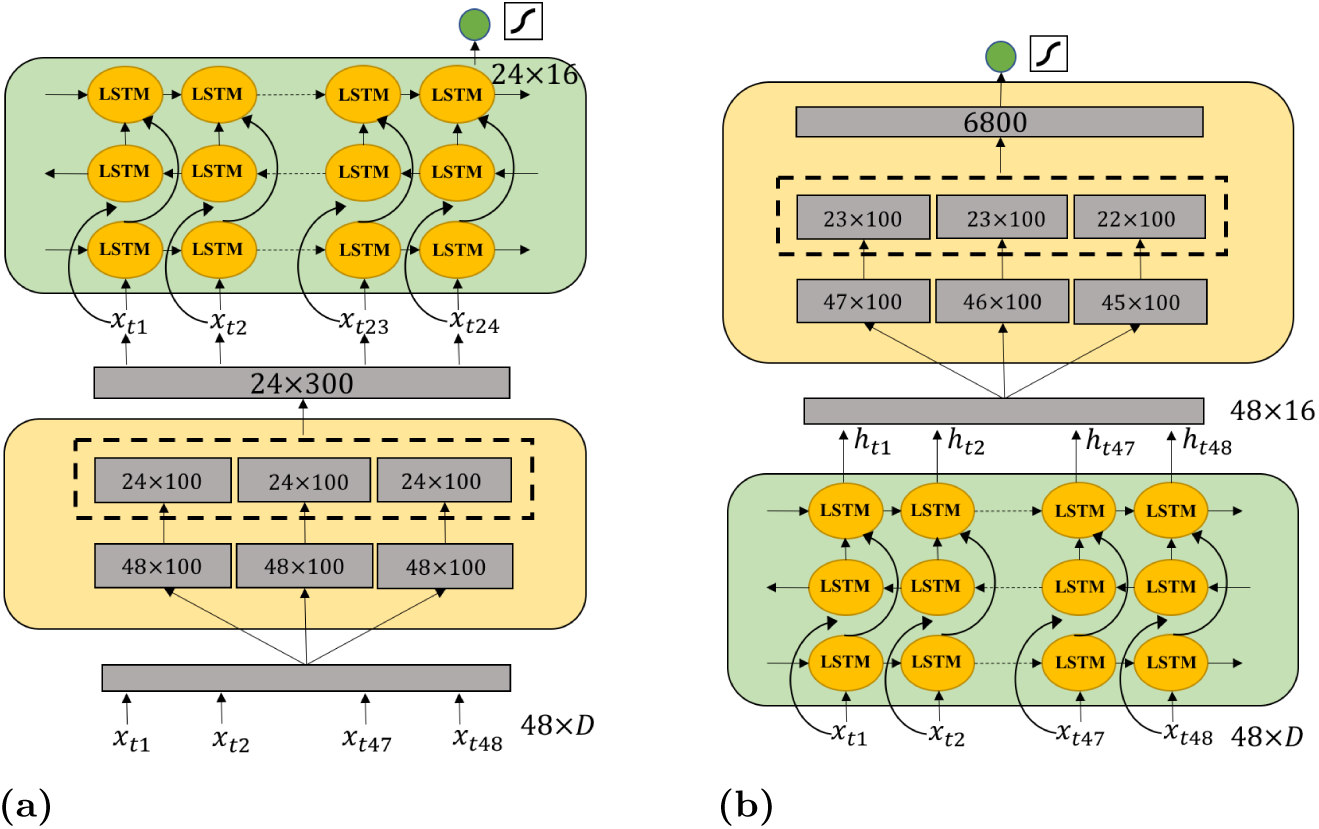
The combination of LSTM and CNN. (a) CNN+LSTM model, and the CNN follows a multi-filter convolution computation with zero padding to maintain the time stamp consistency for different groups of feature maps. The following LSTM only outputs the hidden units of the last time stamp. (b) LSTM+CNN model, and CNN computes the feature maps without zero padding after receiving the output hidden unit sequence from LSTM.

**Fig 4.**
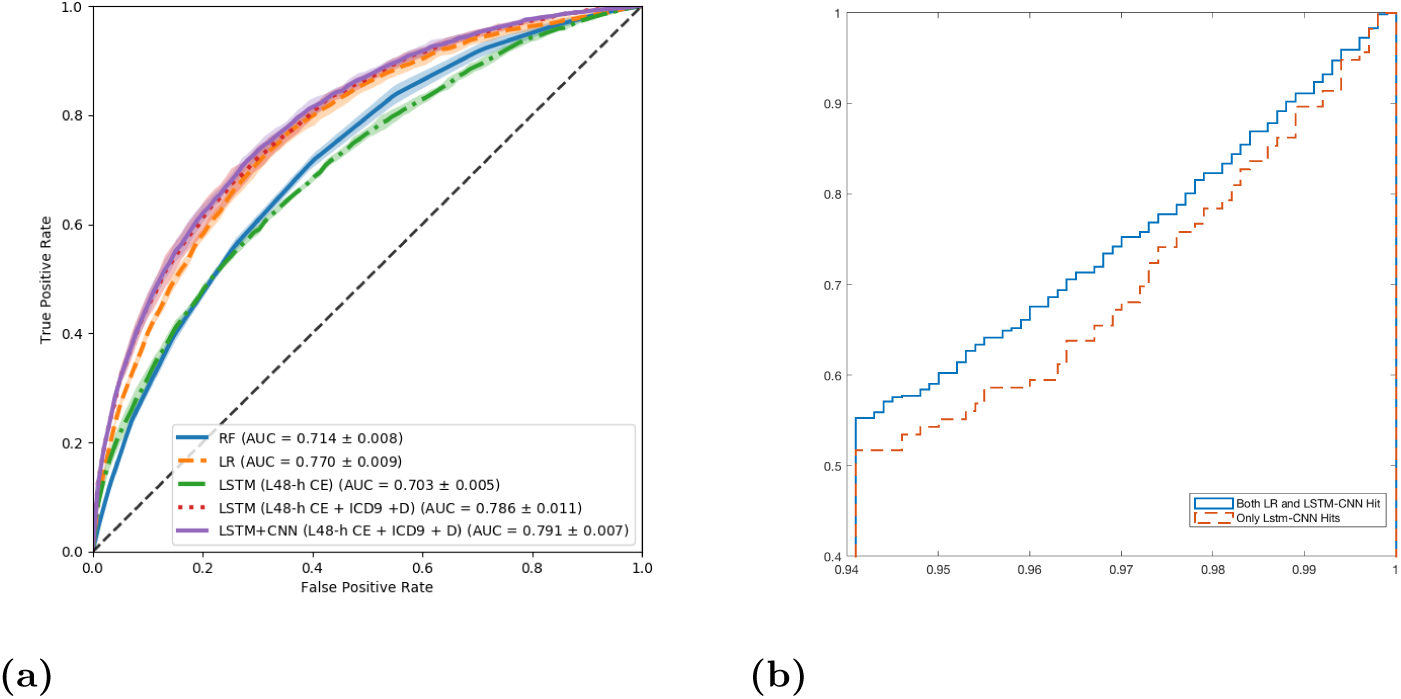
(a) ROC curve of some of the attempted models and features. The color bar is the error bar of the ROC curve with five-fold cross validation. LSTM-CNN model performs relatively better than other ones. Better view in color mode. (b)Cumulative Density Function curve of LSTM-LR-C (blue solid line) and LSTM-C (red dashed line). It suggests that there are higher portions of patient records in the LSTM-C set which have at least one chart event with high oscillation. Therefore, compared to Logistic Regression, the LSTM+CNN model is better for recalling patients with high-oscillated sequence records.

Specifically, CNN model is the 1D multi-filter version introduced in last section, and for the CNN+LSTM model, the CNN follows a multi-filter convolution computation with zero padding to maintain the time stamp consistency for different groups of feature maps. The following LSTM only outputs the hidden units of the last time stamp.

However, for the LSTM+CNN model, CNN computes the feature maps without zero padding after receiving the output hidden unit sequence from LSTM. Our experiment results showed that LSTM followed by a CNN utilizing all the feature sets obtains a higher positive recall rate and overall prediction performance. The proposed model outperforms the traditional approaches trained with statistical features. The ROC curve of some selected models and features are shown in Fig 4a.

## Discussion

In this section, we interpret our model by feature ablation test, and investigate the most important factors the black-box model learns to predict ICU readmission. Finally, we discover the advantages of the proposed model over traditional linear models by studying the statistics of the true positive sets of each model.

### Model Interpretation

We conducted the feature ablation test on the chart events to explain the proposed model. We selected all the positive cases on the testing partition, and obtained all the true positive samples after running the LSTM+CNN model utilizing all the features. These true positive cases are recalled correctly by our proposed model. For each case, every time we changed only one of the chart events to its normal value, and recorded the number of cases to be predicted false. Then we ranked all the chart events according to the numbers and the results are shown in Fig 5. The y-axis is the prediction result changing ratio when we replace the original feature with normal value. It showed that Glucose are the most important factor learned by the black-box model for readmission prediction, while Capillary Refill Rate, Fraction inspired Oxygen, and Systolic Blood Pressure do not influence the prediction result too much. The changing of the prediction is not dramatic, which may result from the correlation among different factors. The corresponding hypothesis could be further validated by back propagation approach which is left for future work.

**Fig 5.**
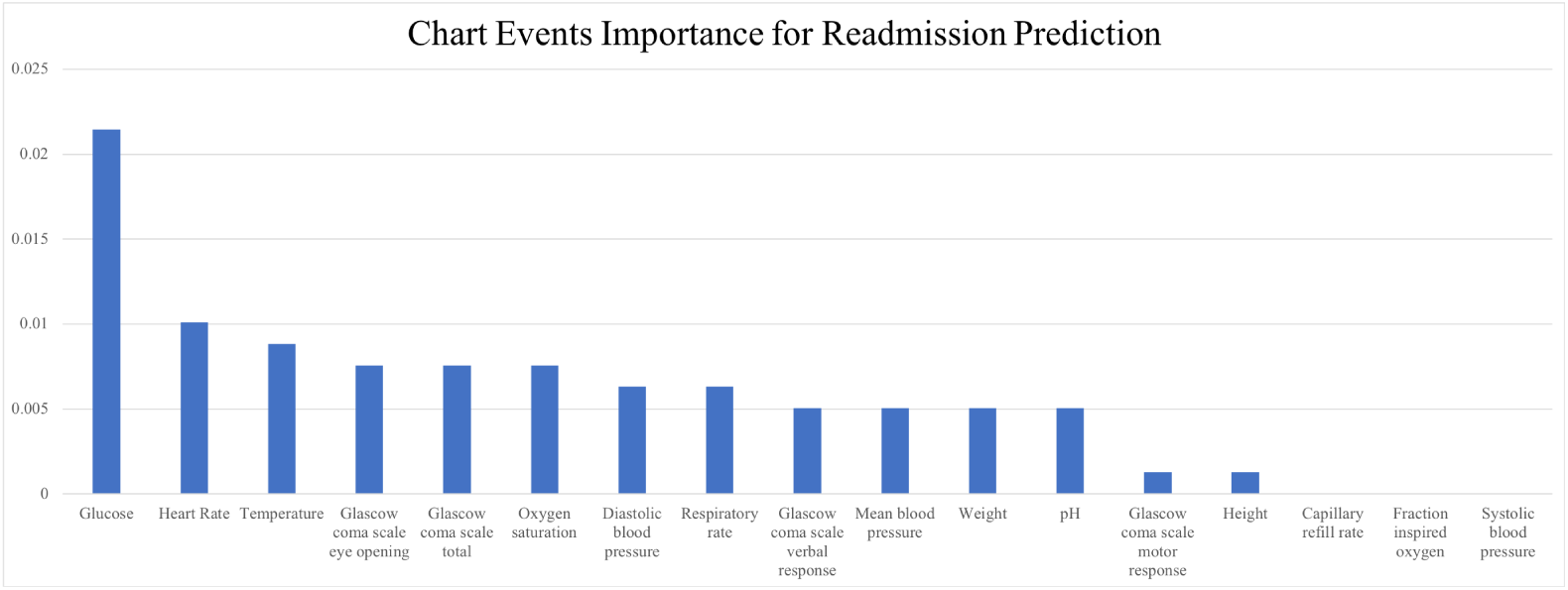
The importance of chart events when predicting the ICU readmission.

### Comparison with Logistic Regression

To verify the advantages of LSTM-based model over the traditional linear model, we investigated the positive patients who are predicted correctly by the LSTM+CNN but misclassified by the logistic regression model. In a randomly selected fold, there are totally 116 positive patients in the testing partition who are only predicted correctly by the LSTM+CNN model. We denote it by LSTM-C set. For another 676 cases predicted correctly by both the LSTM+CNN and Logistic Regression model, we denote the set by LSTM-LR-C.

We measured the degree of value oscillation of each numerical chart event by introducing a factor *D_nm_* called average absolute neighbor difference for record *n* of chart event *m*. Given a numerical chart event sequence *E_nm_* = *{x_t_}*, where *t ∈* [1, 48], then *D*_*nm*_ can be computed by,

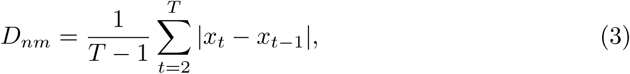

where T equals to the length of one existing record, normally 48 if no missing data.

For each chart event, we picked up the patients with the highest *D*_*nm*_ in the LSTM-C set and plotted all the sequence values of the numerical events of this stay. Two of the examples are illustrated in Fig 6. The common character of these two patients is that the abnormal sequence are oscillated around the normal value of the chart event types. Therefore, linear model will regress it to a normal value with tiny slope, losing important factors of readmission prediction: repeated illness and unstable status.

**Fig 6.**
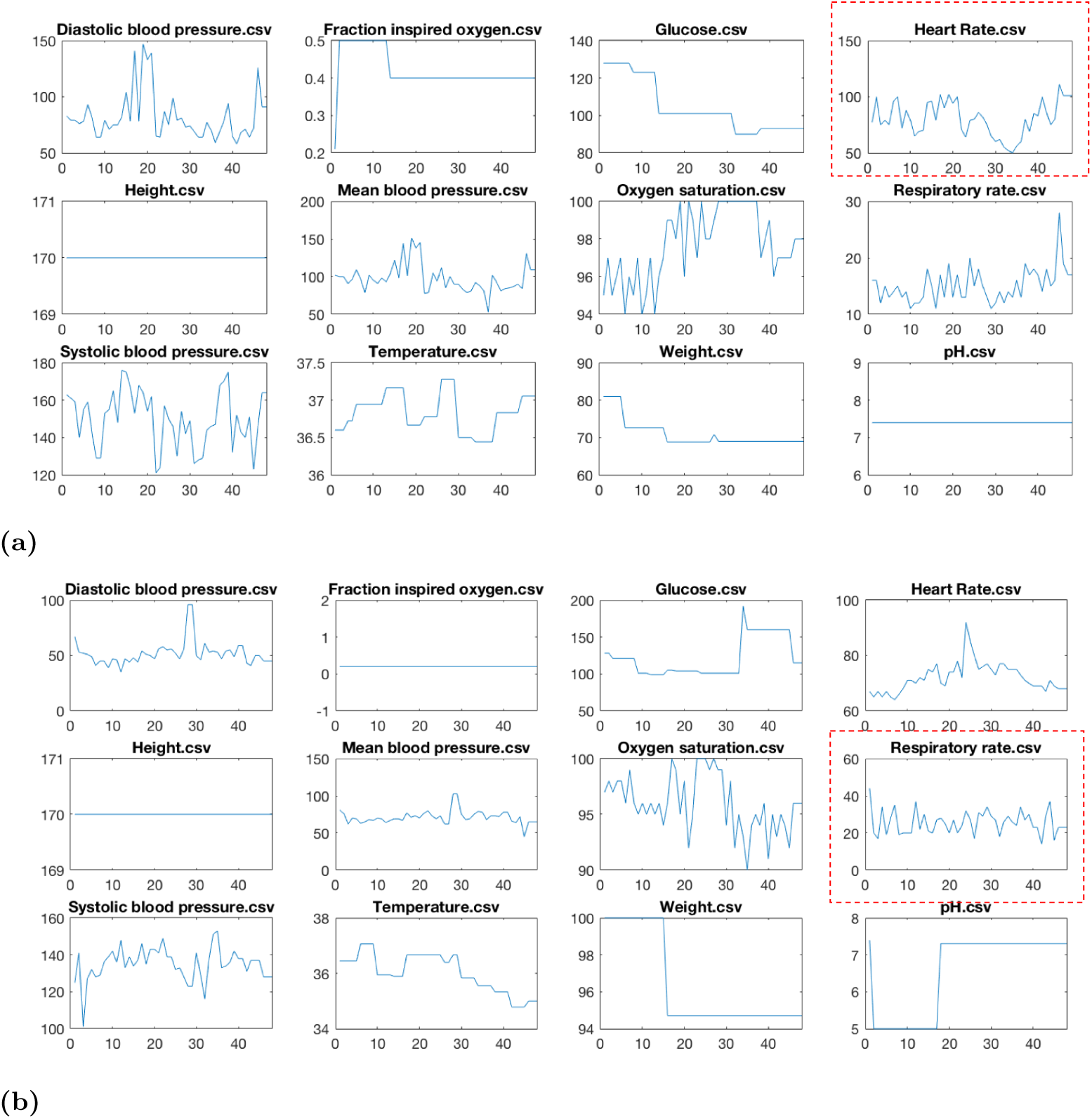
(a)One selected ICU-study sample with the highest *C* of heart rate event, and another sample with the highest *C* of respiration rate. These patients are predicted correctly by the LSTM-CNN model, but wrongly by the traditional linear model. The common character of these two patients is that the abnormal sequences are mostly oscillated around the normal values of the chart event types. Therefore, linear model will regress them to normal value with tiny slopes. It suggests that LSTM can better model the sequence with value oscillation, yielding higher recall rate.

To obtain the big picture of LSTM-C and LSTM-LR-C, we computed the highest oscillation of each stay across all the 12 numerical chart events, and compared the value distributions of the two sets. We first estimated the cumulative density function (CDF) *P*_*m*_ of the histogram of each chart event on the whole positive set. Then we remapped each *D*_*nm*_ to the probability *p*_*nm*_, and computed the maximum probability *w*_*n*_ for each record *n* by,

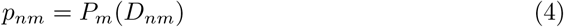

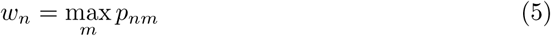

w_n_ represents the highest oscillation among all the chart events for this record.

Finally, for both LSTM-C and LSTM-LR-C set, we plotted the CDFs of the estimated histograms of *w_n_* in Fig 4b. It suggests that there are higher portions of patient records in the LSTM-C set which have at least one chart event with high oscillation. Therefore, compared to Logistic Regression, the LSTM+CNN model is better for recalling patients with high-oscillated sequence records.

## Conclusion

In this study, we addressed the unplanned ICU readmission prediction by utilizing chart events, demographics and ICD9 embeddings features. Among the data that we used, chart event features are significantly sensitive to time series, and cannot be properly captured by traditional static models (e.g., logistic regression). We proposing a LSTM-CNN based model, which can properly incorporate time series data without information lost.

Our model achieved a positive case recall rate (sensitivity) of 0.742, AUROC of 0.791, which contribute to the literatures by improving the sensitivity. Moreover, we illustrated the importance of each portions of the features and different combinations of the models to verify the effectiveness of the proposed model. To further understand the focus of the predictive model, we conducted the chart events ablation test to rank the influence of different factors when predicting the ICU readmission.

In future work, more validation can be conducted on real data of local hospitals and a better way of explaining the deep models should be discovered.

